# Extracellular matrix components are required to protect *Bacillus subtilis* from T6SS-dependent *Pseudomonas* invasion and modulate co-colonization of plants

**DOI:** 10.1101/429001

**Authors:** Carlos Molina-Santiago, John R. Pearson, Yurena Navarro-García, María Victoria Berlanga-Clavero, Andrés Mauricio Caraballo-Rodriguez, Daniel Petras, Francisco M. Cazorla, Antonio de Vicente, Pieter C. Dorrestein, Diego Romero

## Abstract

Bacteria adapt to environmental changes and interact with other microorganisms using a wide array of molecules, metabolic plasticity, secretion systems and the formation of biofilms. Some research has looked at changes in the expression of biofilm related genes during interactions between different bacterial species, however no studies have directly demonstrated the functional significance of biofilms in modulating such interactions. In this study, we have explored this fundamental question by studying the interaction between *Bacillus subtilis* 3610 and *Pseudomonas chlororaphis* PCL1606. We demonstrate the important role of the extracellular matrix in protecting *B. subtilis* colonies from infiltration by *Pseudomonas*. Surprisingly, we find that the *Pseudomonas* type VI secretion system (T6SS) is required in the cell-to-cell contact with matrix-impaired *B. subtilis* cells, revealing a novel role for T6SS against Gram-positive bacteria. In response to *P. chlororaphis* infiltration, we find that *B. subtilis* activates sporulation and expresses motility-related genes. Experiments using plant organs demonstrate the functional importance of these different bacterial strategies in their coexistence as stable bacterial communities. The findings described here further our understanding of the functional role played by biofilms in mediating bacterial social interactions.

## Introduction

Plant-associated microorganisms can be divided into three groups, beneficial^1, 2, 3, 4^, deleterious^5, 6, 7^ and neutral^8, 9^, depending on their effect on plant hosts. Bacteria belonging to these three groups continuously interact with their plant hosts and between themselves, constituting what is known as the plant microbiome^10^. Bacteria found in plant-related environments usually reside in structurally and dynamically complex biological systems known as biofilms, which can confer a range of benefits including increased resistance to environmental stresses, metabolic consortia, better surface colonization, and increased horizontal gene transfer^11, 12^. Common to all bacterial biofilms is the presence of a secreted extracellular matrix that holds the cells together and provides robustness to the biofilm architecture. Although the composition of the extracellular matrix varies between species, in general they are composed of polysaccharides, proteins and nucleic acids^13, 14^.

*Bacillus subtilis* is a soil dwelling bacteria that lives in harmony with plants and is used as a model of biofilm formation^14, 15, 16, 17^. The extracellular matrix of *B. subtilis* is mainly composed of exopolysaccharides, synthesized by the *epsA-O* operon-encoded genes; TasA, a functional amyloid encoded in the three-gene operon *yqxM\tapA-sipW-tasA*^18, 19^; and BslA, which is involved in the formation of a hydrophobic coat over the biofilm^20^. Although their role in biofilm assembly is well understood, little is known about their functional importance in real environments, such as on plant surfaces. Cells growing within a biofilm are in continuous contact with each other, as well as with their environment and other organisms living in the same ecological niche. Bacterial interactions are thus defined by a combination of different factors including the activation of different metabolic pathways in conjunction with the production and secretion of signaling compounds, siderophores, antibiotics and quorum sensing molecules^21, 22, 23, 24, 25, 26^. These compounds can be excreted outside the cells via bacterial secretion systems, efflux pumps, transporters and membrane vesicles. Secretion mechanisms can be compound-specific or be able to extrude a broad spectrum of compounds, with some requiring tight physical connections between the interacting cells, such as the type VI secretion system (T6SS), a tubular puncturing device able to deliver molecules directly into other cells. For example, bacteria of the genus *Pseudomonas*, can use their T6SS to manipulate and subvert eukaryotic host cells and/or to fight other bacteria thriving in the same ecological niche^27,28^.

*Pseudomonas* spp. and *Bacillus* spp. are among the most predominant genera of plant-beneficial bacteria. Both genera have been well studied and their abilities to protect plants against pathogens^29, 30, 31^ and to promote growth of many plant species have been widely described^32, 33^. However, studies examining how these bacterial species interact and cohabit are scarce, and the limited reports on the antagonistic relationship between the two species were addressed by *in vitro* experiments^34, 35^. It has been shown that the genetic pathway that regulates *B. subtilis* biofilm assembly is altered in these interactions, but the mechanistic basis by which changes to the extracellular matrix affect bacterial interactions remains unknown^35^.

In this work, we explore the functional role of the *B. subtilis* NCIB3610 extracellular matrix in its interaction with *Pseudomonas chlororaphis* PCL1606, a strain isolated from the plant rhizosphere possessing antifungal activity^1, 36, 37^. We find that the extracellular matrix represents a vital defense against invasion by *P. chlororaphis*. Time-lapse confocal microscopy reveals differences in the expansion rates of the bacterial colonies and allowed us to follow, at the cellular level, how *Pseudomonas* can infiltrate *Bacillus* colonies when they lack an extracellular matrix. RNA-seq analysis identified that *Pseudomonas* T6SS genes are triggered by contact with *Bacillus*. We also identify the expression of *B. subtilis* genes involved in motility and the activation of sporulation as responses to *Pseudomonas* invasion. Imaging metabolomic analysis of interacting bacteria suggests that altered expression of the lipopeptide surfactin may have a role facilitating the spread of *P. chlororaphis*, thus influencing initial contact between the two bacterial species. Analysis of co-bacterizations on plants support the key role for the *Bacillus* extracellular matrix in determining bacterial distribution in mixed populations on leaves and a determinant role for the *Pseudomonas* T6SS during plant seeds germination.

## Results

### The extracellular matrix of *Bacillus subtilis* NCIB3610 has a protective role in the interaction with *P. chlororaphis* PCL1606

Previous studies have reported the differential expression of extracellular matrix components in interactions between *Bacillus* and other bacterial species^35^. These findings supported a hypothetical contribution of the extracellular matrix to the adaptation of *Bacillus* to the presence of other bacterial species, but no studies have directly demonstrated the functional significance of this bacterial structure in modulating such interactions. Therefore, to better understand the function of the extracellular matrix in such exchanges, we decided to study the interaction between *B. subtilis* NCIB3610 (referred to as 3610) and *Pseudomonas chlororaphis* PCL1606 (referred to as PCL1606), two widely distributed bacterial species that are highly likely to co-exist on plant surfaces. We initially evaluated the behavior of these strains with pairwise interaction experiments using four different artificial media: King’s B, a medium optimum for the growth and production of secondary metabolites of *Pseudomonas* strains; Msgg a medium optimum for the study of biofilm formation in *Bacillus*; and TY or LB, two rich media routinely used for the growth of organotrophic bacterial species (Suppl. Fig. 1). Interactions after 72 h of growth showed the existence of a clear halo separating PCL1606 and 3610 colonies, an effect especially evident in King’s B medium; and a reduction of wrinkles in the 3610 colony morphology. From this preliminary analysis, we decided to use LB medium to investigate the mechanism behind this bacterial interaction for two main reasons: first, the similar growth rate of both strains in LB, thus permitting a “balanced” interaction; and second, the apparent differences in *B. subtilis* biofilm morphologies.

We built single, double and triple mutants of all the *B. subtilis* extracellular matrix components to investigate their respective contribution to the interaction with PCL1606. Pairwise time-course interactions between PCL1606 and WT 3610 revealed the existence of a subtle inhibition area between the two colonies, and especially the reduction of wrinkles in 3610 versus the strain growing alone (Fig. 1a and d). Interactions with single mutants in *tasA* and the double mutant in *tasA* and *bslA* were similar to those obtained for WT 3610 (Suppl. Fig. 2a, b and d). However, in single mutants for Eps, and to a lesser extent for BslA, PCL1606 was able to penetrate the *B. subtilis* colony after 72 h (Suppl. Fig. 2c and f), and to partially colonize the frontline of the *Bacillus* colony after 96 h. This behavior was even more evident in the interaction between PCL1606 and the triple *eps, tasA, bsla* mutant (referred to as Δmatrix) where PCL1606 was able to completely colonize the *B. subtilis* colony after 120 h of interaction (Fig. 1b and e). These findings strongly suggest a defensive role for the extracellular matrix, and that Eps and BslA are particularly important for this interaction.

**Fig. 1.**
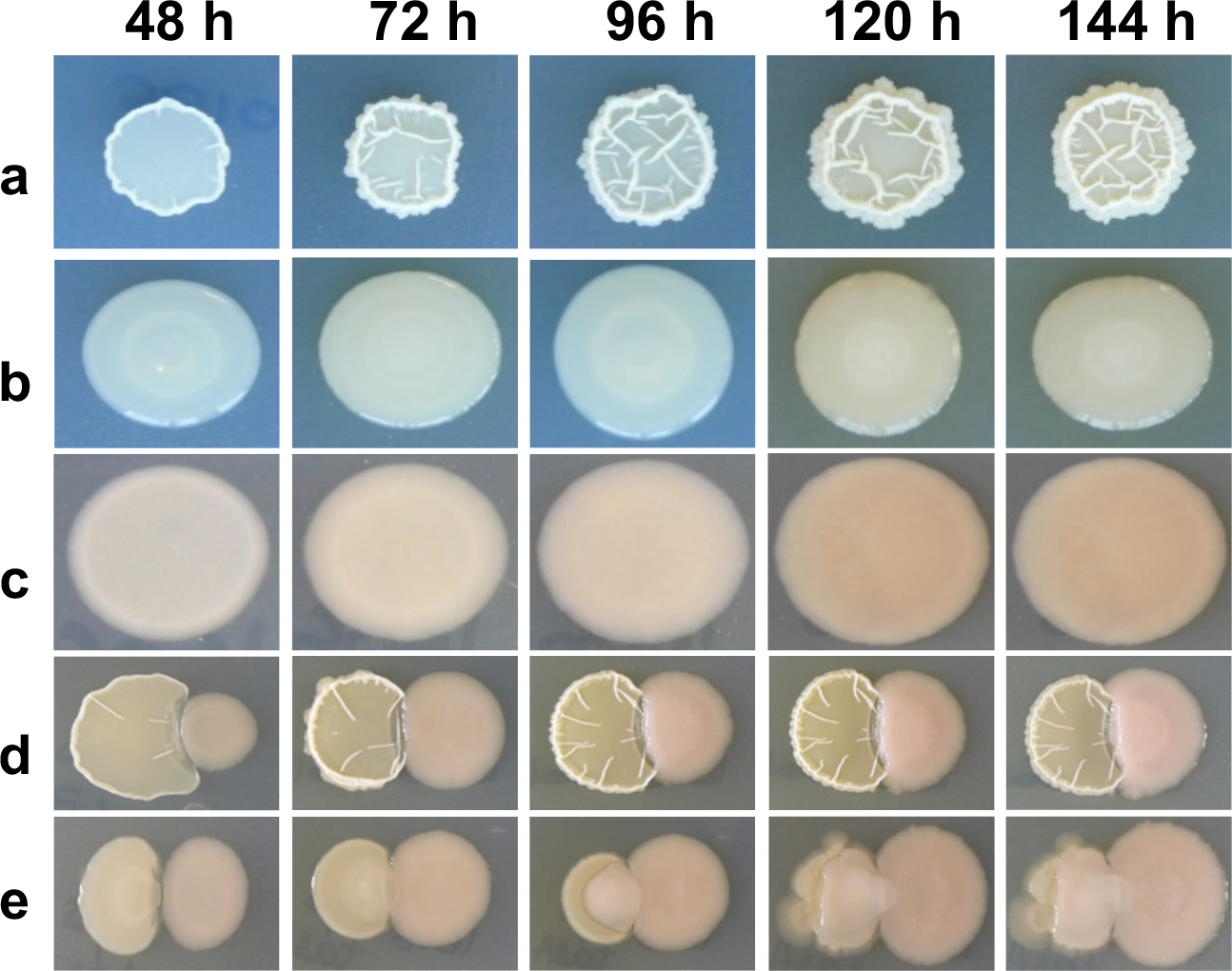
The lack of *B. subtilis* extracellular matrix permits the overgrowth of *P. chlororaphis*. (a, b and c) Time-courses of *B. subtilis* (a) 3610, (b) Δmatrix and (c) *P. chlororaphis* PCL1606 colony morphologies at 48 h, 72 h, 96 h, 120 h and 144 h when growing alone in LB medium. (d and e) Time-courses of the interactions (d) 3610-PCL1606 and (e) Δmatrix-PCL1606. In the interactions, left colonies are *B. subtilis* strains and right colonies are *P. chlororaphis*.

### *P. chlororaphis* PCL1606 covers but does not induce widespread death of *B. subtilis* cells lacking extracellular matrix

We had seen that the PCL1606-Δmatrix interaction culminates with PCL1606 penetrating the Δmatrix colony. We hypothesized that *B. subtilis* cells were being killed after physical contact between the colonies. To test this idea, we measured the colony forming units (CFUs) for each species in three different areas of the interaction: a *Pseudomonas* area corresponding to the initial zone where the *Pseudomonas* colony was spotted (Fig. 2-I and Suppl. Fig. 3-I), an intermediate area corresponding to the zone where the first contact between the two colonies took place (Fig. 2-II and Suppl. Fig. 3-II), and a *Bacillus* area corresponding to the zone where the *Bacillus* colony was initially spotted (Fig. 2-III and Suppl. Fig 3-III). As expected from our macroscopic observations, no *B. subtilis* CFUs (either 3610 or Δmatrix) were obtained in the *Pseudomonas* area at any time during the interaction (Fig. 2a and 2d; and Suppl. Fig. 3a and 3d). The intermediate area was dominated by PCL1606 with percentages up to 95% of the total population (Fig. 2b and Suppl. Fig. 3b). However, and contrary to our expectations based on macroscopic observations, *Bacillus* CFUs were obtained at all time-points, and no differences in population size were evident between early (24 h) and later (96 h) stages of the experiment (Fig. 2e and Suppl. Fig. 3e). The most noticeable differences were observed in the *Bacillus* area. In the interaction between the two wild-type strains, *B. subtilis* CFUs made up virtually all the bacteria detected with PCL1606 CFUs representing 0.007% of the total observed at later stages of the interaction (96 h), confirming that PCL1606 cannot easily penetrate 3610 colonies with a functional extracellular matrix (Suppl. Fig. 3c and 3f). As expected, in the Δmatrix-PCL1606 interaction, a higher PCL1606 population was detected in the *Bacillus* area after 48 h, and progressively increased over time becoming the majority at 72 and 96 h (Fig. 2c). Surprisingly, however, the *Bacillus* CFUs remained unchanged during the time-course experiment, even after contact with PCL1606 (Fig. 2e). One possibility is that invasion might trigger a survival mechanism in *Bacillus* bacteria. Indeed, the sporulation rates of 3610 and Δmatrix bacteria increased dramatically from 58% and 30%, respectively, in monocultures (Suppl. Fig. 4a and 4b) to 65% in Δmatrix and 95% in 3610 wild-type strain, upon initial contact with PCL1606 after 48 hours in the intermediate area (Fig. 2b, Suppl. Fig. 3b and Suppl. Fig. 4a and b) and in the case of Δmatrix, reaching 95% in the *Bacillus* area after 72 h.

**Fig. 2.**
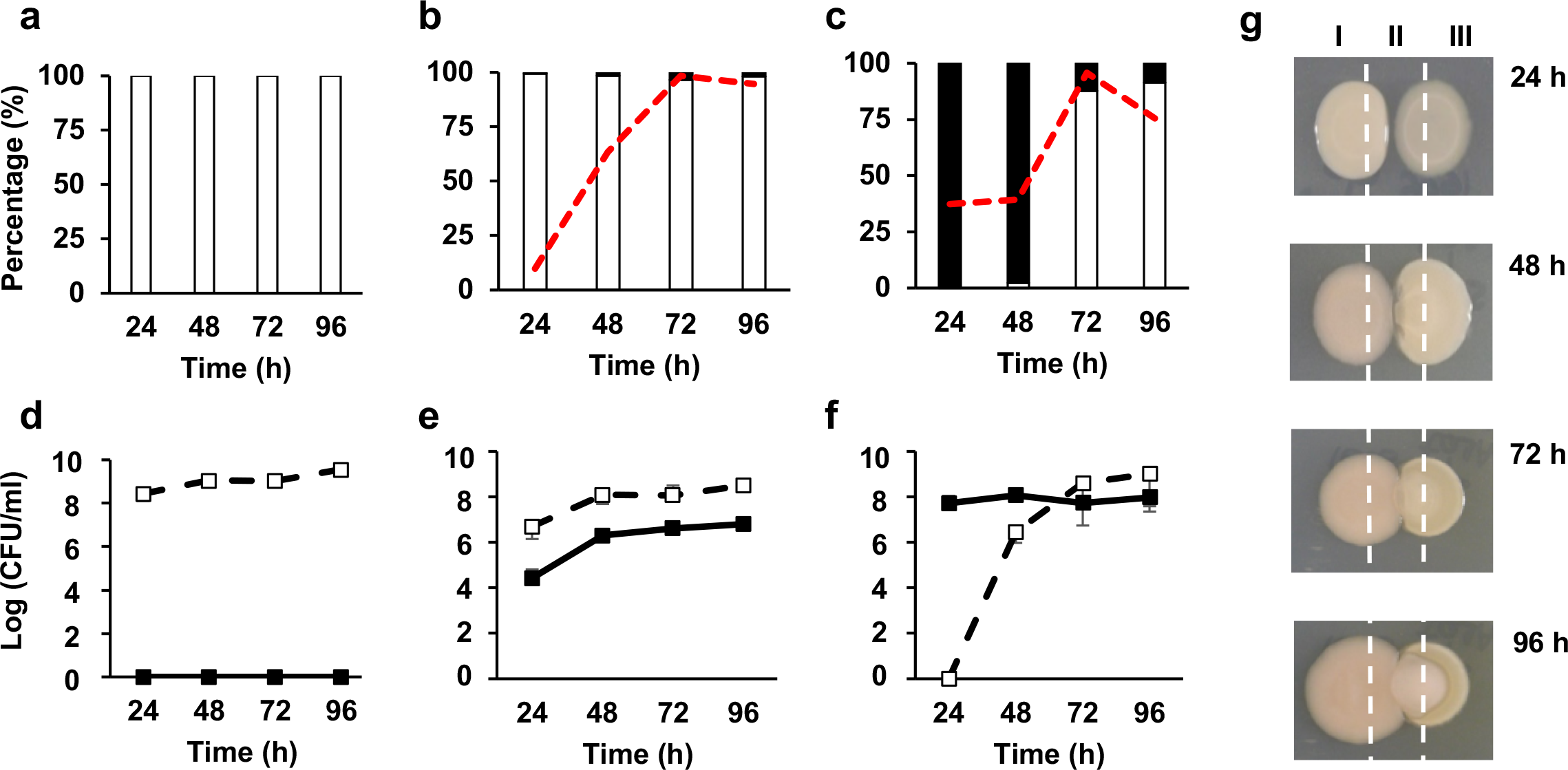
B. subtilis Δmatrix coexists in the form of spores with P. chlororaphis PCL1606. PCL1606 is shown as (a, b, and c) empty bars or (d, e, and f) empty squares; Δmatrix is shown as (a, b, and c) black bars or (d, e, and f) black squares. Sporulation rates are shown as red lines. (a, b and c) Percentage and (d, e and f) Log (CFU/ml) of PCL1606 and Δmatrix colony forming units and sporulation rates at each stage of the interaction (24 h, 48 h, 72 h, 96 h). (g) Interactions were divided in three sections: (I) PCL1606 area, (II) Intermediate area and (III) Δmatrix area. Cells from the three sections of the interaction were collected, diluted and plated to determine colony forming units (CFU), sporulation rates and percentages of each species in the interaction. Average values of three biological replicates are shown, with error bars representing SD.

Taken in combination, these results support the importance of sporulation as a secondary protective mechanism for *B. subtilis* in the presence of a competitor, such as PCL1606, thus permitting the strain to survive even though it lacks an extracellular matrix.

### The initial stages of the Δmatrix - PCL1606 interaction are dominated by an aggressive expansion of the *B. subtilis* colony

To better understand the infiltration of Δmatrix colonies by PCL1606 cells, we decided to study this interaction at the cellular level using fluorescently-labeled strains and time-lapse confocal laser scanning microscopy (CLSM). A PCL1606 strain constitutively-expressing DsRed and CFP-expressing *B. subtilis* strains were spotted at a distance of 0.7 cm from each other on LB agar medium and their growth and interactions followed by time-lapse microscopy. Images were captured every 30 minutes over a total period of 4 days at multiple positions on the plate for each time-point. This procedure allowed us to define different stages of the interaction: independent colony growth, indirect interactions, first direct contact, and colony interaction resolution (Fig. 3). Furthermore, analysis of each time-lapse video allows colony expansion rates to be calculated for each strain over the course of the interaction process.

**Fig. 3.**
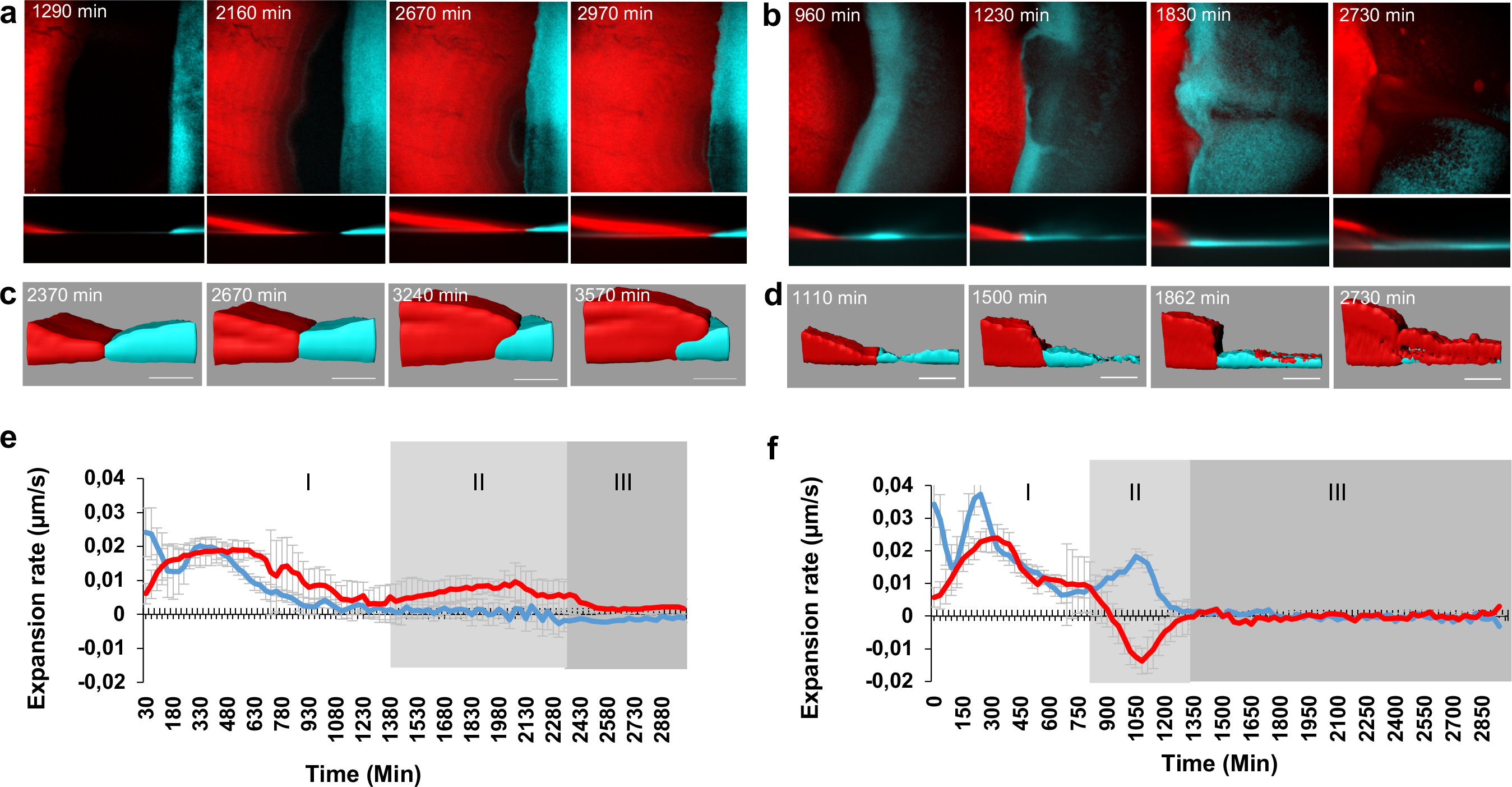
The expansion rate of the colonies changes during the interaction. Red lines represent PCL1606 and blue lines indicate *Bacillus* strains. (a, b, c, d) CLSM time-course experiments of the interactions between (a and c) PCL1606 and 3610 and (b and d) PCL1606 with Δmatrix. (a) and (b) show the main steps during the interactions as a sum of the projections and in z-axis. (c) and (d) show the most important points since both bacteria have interacted. (e and f) Expansion rates of the *Bacillus* and *P. chlororaphis* strains during the interactions can be divided in three different stages (I, II, III). (e) shows the expansion rates of 3610 and PCL1606 during the interaction. (f) shows the expansion rates of Δmatrix and PCL1606 during the interaction. Average values of four biological replicates are shown, with error bars representing SD.

Microscopic observations of the interaction between wild-type 3610 and PCL1606 were in line with our expectations based on the macroscopic analysis described above. During the first hours after being spotted onto the culture medium, wild-type 3610 cells started growing with a spatial distribution similar to the previously described Van Gogh bundles^38^, with phases of advancement followed by phases of filling the colonized area (Fig. 3a and Suppl. Movie 1). During this period of time (first 18-20 h), PCL1606 expanded, on average, faster than 3610 (0.013 µm/s vs 0.010 µm/s, respectively) (Fig. 3e-I). As the colonies approached, 3610 stopped expanding while PCL1606 continued spreading, however, the speed of the expansion progressively decreased until the colonies finally contacted after 30-35 hours of growth (Fig. 3a, c and e-II). From this time on, a subtle and slow growth of PCL1606 over 3610 in the interaction zone was observed (Fig. 3a, c and e-III). Thus, under normal circumstances the two colonies retain their integrity and general organization, with a clearly defined boundary that changes only gradually.

Time-lapse imaging of Δmatrix and PCL1606 interactions uncovered a very different interaction, providing some remarkable and unexpected insights into the process (Fig. 3b and Suppl. Movie 2). First, Δmatrix cells expanded significantly faster (0.017 μm/s) than the wild-type 3610 and PCL1606 (0.013 μm/s) strains during the first 10-16 h of the experiment (Fig. 3f-I), which may indicate higher liquidity compared to wild-type 3610 colonies. Second, at the moment of first direct contact at 20-24 hours, the speed of Δmatrix expansion unexpectedly increased and started to grow over the PCL1606 colony surface, an interaction that is macroscopically imperceptible (Fig. 3b and d). This critical stage can also be described by measuring the expansion rate of PCL1606, with negative values indicating a regression in the position of this strain on the solid medium (Fig. 3f-II). Third, when the interaction seemed to have stabilized after a few hours, CFP-labeled Δmatrix cells started disappearing from the interaction zone, presumably due to sporulation of the population (Fig. 3b). In parallel, we observed a concomitant emergence of *Pseudomonas* spots-colonies, which resolved with the complete colonization of the Δmatrix colony (Fig. 3b and d). Due to almost imperceptible nature of the early stages of the invasion, presumably by individual *Pseudomonas* cells, it was not possible to accurately calculate speed measurements at this stage (Fig. 3f-III). Thus, the loss of extracellular matrix results in increased colony expansion speed but at the cost of reduced resistance to invasive bacteria.

### Differences in the spatial distribution of *B. subtilis* surfactins suggest a role in bacterial interactions and motility

The altered kinetics of Δmatrix and PCL1606 colony interactions could be interpreted primarily as a change in the biophysical properties of the Δmatrix colony biofilm that is then exploited by *Pseudomonas*. However, some observations suggest that the situation may be more complex. For example, the acceleration of Δmatrix and PCL1606 cells observed by CLSM at the earlier stages of their interaction might indicate the presence of molecules that favor bacterial cell movement. To clarify this moment of the interaction, we initially studied the spatial distribution of metabolites by performing imaging mass spectrometry analyses of single colonies and interactions at 24 h, 48 h and 72 h (Figure 4a, b, c and d). We did not see significant changes to the composition or spatial distribution of *Pseudomonas* metabolites between single colonies and wild-type and Δmatrix interactions (data not shown). In general, most metabolites produced by *B. subtilis* showed similar distribution patterns in both interactions and monocultures, with the exception of surfactin, which is not produced by PCL1606 (Suppl. Fig. 5). A similar composition of surfactin isoforms was observed in the interactions of 3610 or Δmatrix with PCL1606, with the C13, C14, and C15-surfactin isoforms being the most abundant (Fig. 4e), although their distribution patterns were different. In individual wild-type 3610 colonies, surfactin mainly accumulated outside of the colony at 24 and 48 hours before decreasing after 72 hours (Fig. 4a). Significantly, in single Δmatrix colonies the distribution of surfactin was altered with similar levels present both inside and outside the colony, with the biggest differences at 24 and 48 hours (Fig 4b).

**Fig. 4.**
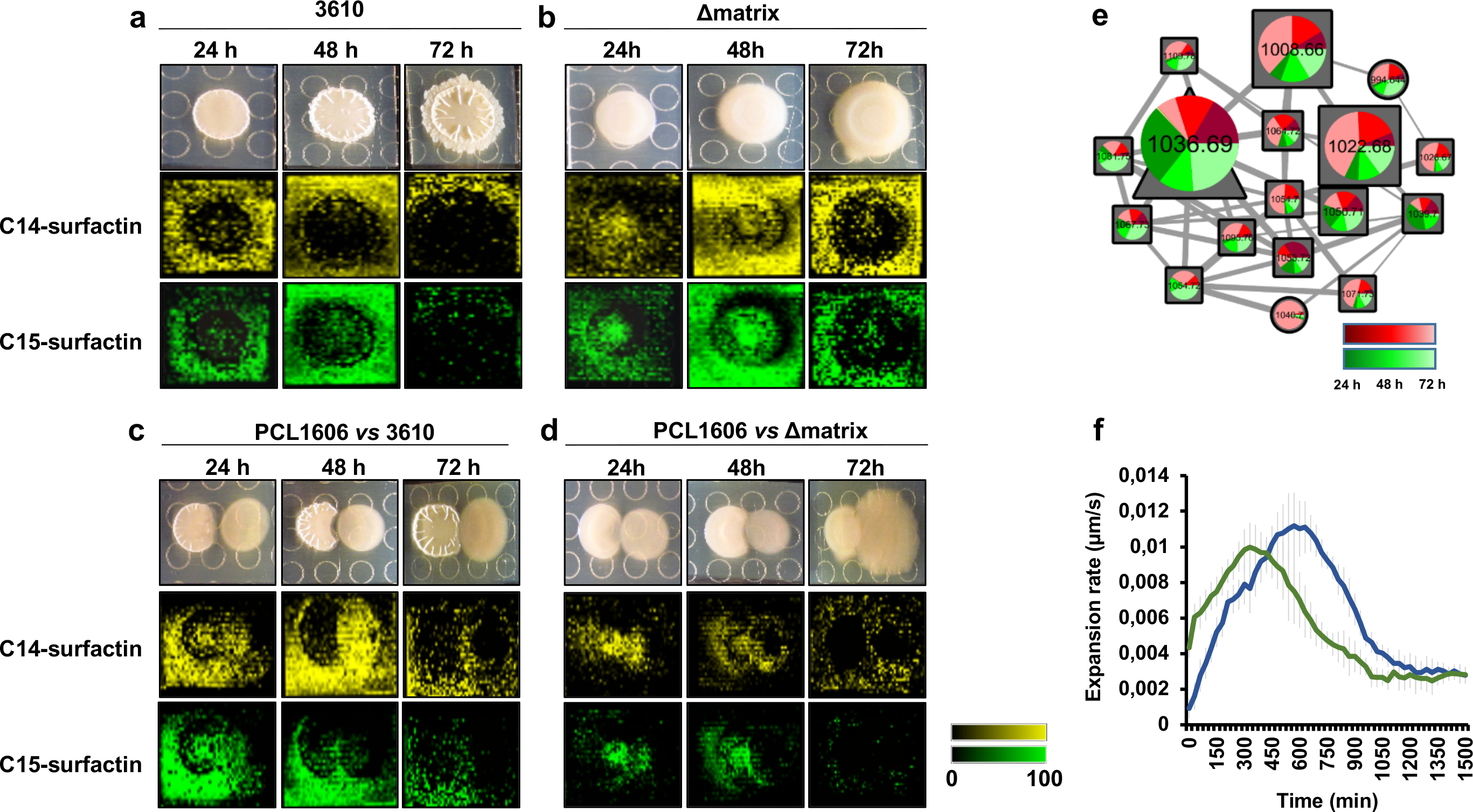
Surfactins produced by *Bacillus* are spatially distributed between Δmatrix and PCL1606 colonies and accelerates the spread of PCL1606. (a, b, c and d) MALDI-IMS time-lapse experiments (24 h, 48 h, 72 h) showed the spatial distribution of C14 and C15-surfactin isoforms (m/z 1058 and 1076 M+Na^+^) when growing (a) 3610 and (b) Δmatrix as single-colonies or during the interactions between (c) PCL1606 (right colonies) and 3610 (left colonies) or (d) PCL1606 (right colonies) and Δmatrix (left colonies). (e) View of the molecular networking results of the surfactins cluster (M+H^+^). Red portions indicate surfactins in Δmatrix single-colonies while green portions indicate surfactins in the interaction Δmatrix-PCL1606 at 24 h, 48 h and 72 h. Size is directly related with abundance of the molecules. C13, C14 and C15-surfactin isoforms correspond to m/z 1008, 1022 and 1036 (M+H^+^), respectively. (f) Expansion rates of PCL1606 in the absence (blue line) or the presence (green line) of surfactin (1 mg/ml) spotted in a sterile disk (n=3).

When we analyzed the distribution of surfactins during the PCL1606-Δmatrix interaction, we surprisingly found that they mostly localized to the interaction area, but were also present in the Δmatrix and PCL1606 colony zones at 24 h and 48 h (Fig. 4d). In contrast, surfactin from wild-type 3610 colonies during their interaction with PCL1606 accumulated strongly around the outside of the colony and throughout the PCL1606 colony but was largely excluded from the *Bacillus* colony itself (Fig. 4c).

These findings could reflect a role for surfactin as a factor that promotes *Bacillus* population motility and thus contributing to the penetration of *Bacillus* Δmatrix cells in the frontline of PCL1606 at early stages of the interaction, as visualized by microscopy (Fig. 3b and d). At the same time, an intriguing possibility is that PCL1606 may hijack surfactin to accelerate its motility and further infiltrate *Bacillus* colonies deprived of an extracellular matrix. In agreement with this possibility, we found that DsRed-labeled PCL1606 colonies expanded with altered kinetics when cultured in the presence of a disk impregnated with 1 mg/ml of surfactin compared to buffer (Fig. 4f). In the presence of surfactin, colony expansion was significantly faster during the earlier stages of the experiment. This effect appears to be relatively short term as control colonies were later able to expand at least as fast but only after a ~5 hours delay. A possible explanation is that exogenous surfactin aids PCL1606 motility in the short term but that limiting factors, such as the rate of bacterial proliferation, may prevent a more sustained increase in the rate of colony expansion.

### *Pseudomonas* activates the T6SS when in close contact with *Bacillus* cells lacking extracellular matrix

So far our results suggest that *Pseudomonas* takes advantage of a deficient extracellular matrix and structural weakness, possibly aided by the altered distribution of surfactin, to enter deeply into the colony, at which point most *Bacillus* undergo sporulation. Next we wanted to use our *in vitro* model to determine which mechanisms were being used by *Pseudomonas* to colonize Δmatrix colonies. We found that *Pseudomonas* mutants unable to produce well-known secondary metabolite did not arrest the overgrowth phenotype (Suppl. Fig. 6). In an effort to identify other pathways and factors that might be involved in this process, we performed a dual RNA-seq analysis looking for changes in gene expression between individual colonies and the Δmatrix-PCL1606 interaction zone at 72 h. Mapping of the sequenced mRNAs to reference genomes showed that the vast majority of hits belonged to PCL1606, consistent with its larger CFU counts and the high sporulation rate of *Bacillus* in the interaction. Analysis revealed 318 differentially expressed genes in PCL1606 during the overgrowth (72 h of interaction), of which 62% were upregulated versus PCL1606 alone (Suppl. Fig. 7a).

The main transcriptional changes in PCL1606 can be categorized into three main groups: i) amino acid and carbon metabolism, ii) membrane composition, and iii) the activation and repression of ABC transporters, efflux pumps and signaling molecules (acyl-homoserine lactones, and siderophores such as pyochelin) (Fig. 5a, Suppl. Fig. 7b and Suppl. Table 1). Among the differentially expressed PCL1606 genes, we also found to be induced genes involved in the formation of the type VI secretion system (T6SS), which is known as an important mechanism in interactions and pathogenesis against bacterial and eukaryotic cells^27,28^, but no studies have reported a role for this system against Gram-positive bacteria. Based on that, we decided to study the role of the T6SS in the interaction between *Pseudomonas* and *Bacillus* cells at a cellular level. For that, the T6SS promoter was fused to the DsRed gene and introduced into PCL1606, and this strain was tested in pairwise interactions against CFP labeled *Bacillus* strains. CLSM analysis revealed a high basal level of the *P*_T6SS_ expression in the PCL1606 colony growing alone after 72 h (Fig. 5b, c and d). In agreement with the RNA-seq expression data, increased fluorescence signal accumulated in locations where PCL1606 and Δmatrix cells were in close contact (Fig. 5e, f and g).

**Fig. 5.**
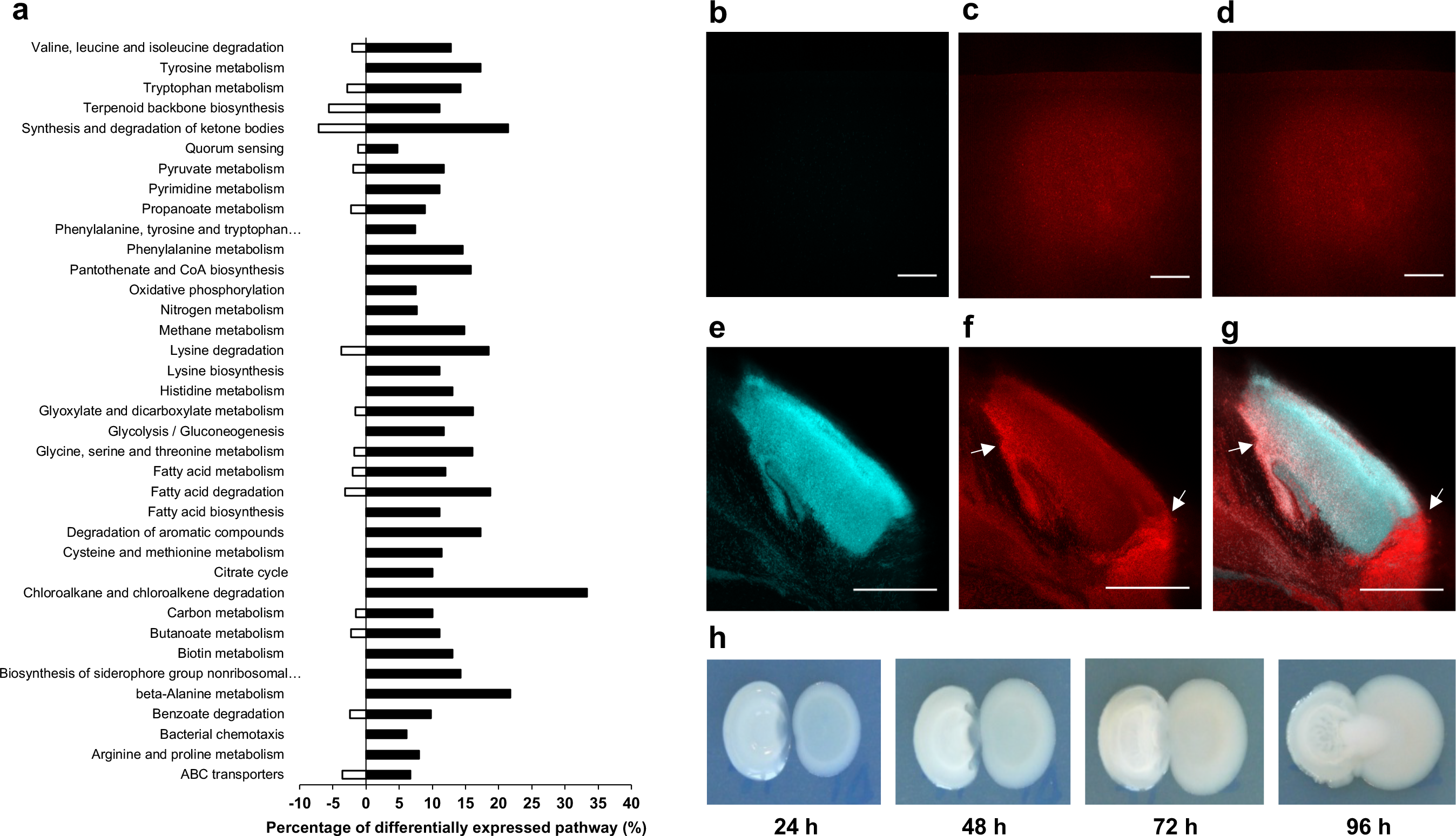
PCL1606 activates the synthesis of the T6SS in the tight contact with *B. subtilis* Δmatrix. (a) KEGG pathway analysis of the PCL1606 genes induced (black bars) and repressed (empty bars) after 72 h of interaction with Δmatrix. (b, c, d, e, f and g) Confocal scanning laser microscopy images of cross-sections of (b, c and d) PCL1606 *P*_*T6SS*_-DsRed colony and (e, f and g) the interaction PCL1606 *P*_*T6SS*_-DsRed *vs* Δmatrix after 72 h of growth. (b and e) Fluorescence from the CFP channel is shown in cyan in images from the CFP channel. (c and f) Fluorescence from the RFP channel is shown in red in images from the RFP channel and (d and g) in the merged images. The fluorescence of the *P*_*T6SS*_-DsRed is shown in red. White arrows indicate the areas where *P*_*T6SS*_ is induced. Scale bar, 100 μm. (h) Time-course images of the interaction ΔT6SS-Δmatrix. Left colonies are Δmatrix strain and right colonies are ΔT6SS.

To better determine the functionality of the T6SS for the invasiveness of *Pseudomonas*, we mutated the *tssA* gene (strain referred to as ΔT6SS), a key component of the T6SS baseplate and critical for the assembly and functionality of the T6SS^29^. Time-course interactions, CFU counts and sporulation percentages of Δmatrix in the interaction with ΔT6SS mimicked the results obtained in the interaction with PCL1606 wild-type strain, including the same pattern of infiltration and colonization of *Pseudomonas* cells on Δmatrix colonies (Fig. 5h and Suppl. Fig. 8). These results highlight the importance of the T6SS in the cell-to-cell contact with Δmatrix although the invasiveness cannot be associated only to this system.

### The remaining vegetative *B. subtilis* Δmatrix cells activate a runaway response in the close contact with *Pseudomonas*

Our previous observations showed that, after 72 h of interaction with PCL1606, around 5% of the Δmatrix population would remain in the form of vegetative cells (Fig. 2c). Therefore, we expected the majority of the differentially-expressed *Bacillus* genes in the RNA-seq would correspond to this 5%. This idea was supported by complementary observations: i) at 72 h almost the entire population of Δmatrix had sporulated; ii) microscopic observations revealed spores released from the mother cells (Suppl. Fig. 9) and iii) the low efficiency of RNA purification from spores using standard protocols^39, 40^. As described above for PCL1606, we compared gene expression between single Δmatrix colonies and *Bacillus* cells at the interaction zone after 72 h, detecting 1105 differentially expressed genes of which 43% were upregulated (Suppl. Fig. 7a).

In contrast to the offensive strategy employed by *Pseudomonas*, the 5% of the *Bacillus* population that remained in a vegetative state arrested the growth, energy consumption and secondary metabolism together with the down-regulation of sporulation and biofilm pathways where the spo0A master regulator is involved (Fig. 6a). On the other hand, Δmatrix cells in the interaction zone activated the machinery of synthesis and reparation of DNA, the PBSX and SPβ prophages, chemotaxis and flagellar assembly, sulfur uptake and nitrogen metabolism, and competence genes such as *comG* and *comE* among other functions (Suppl. Table 2, Fig. 6a and Suppl. Fig. 10). This transcriptomic response appears indicative of attempts by non-sporulating cells to survive the PCL1606 colonization by arresting their growth while activating escape strategies represented by the chemotaxis and motility.

**Fig. 6.**
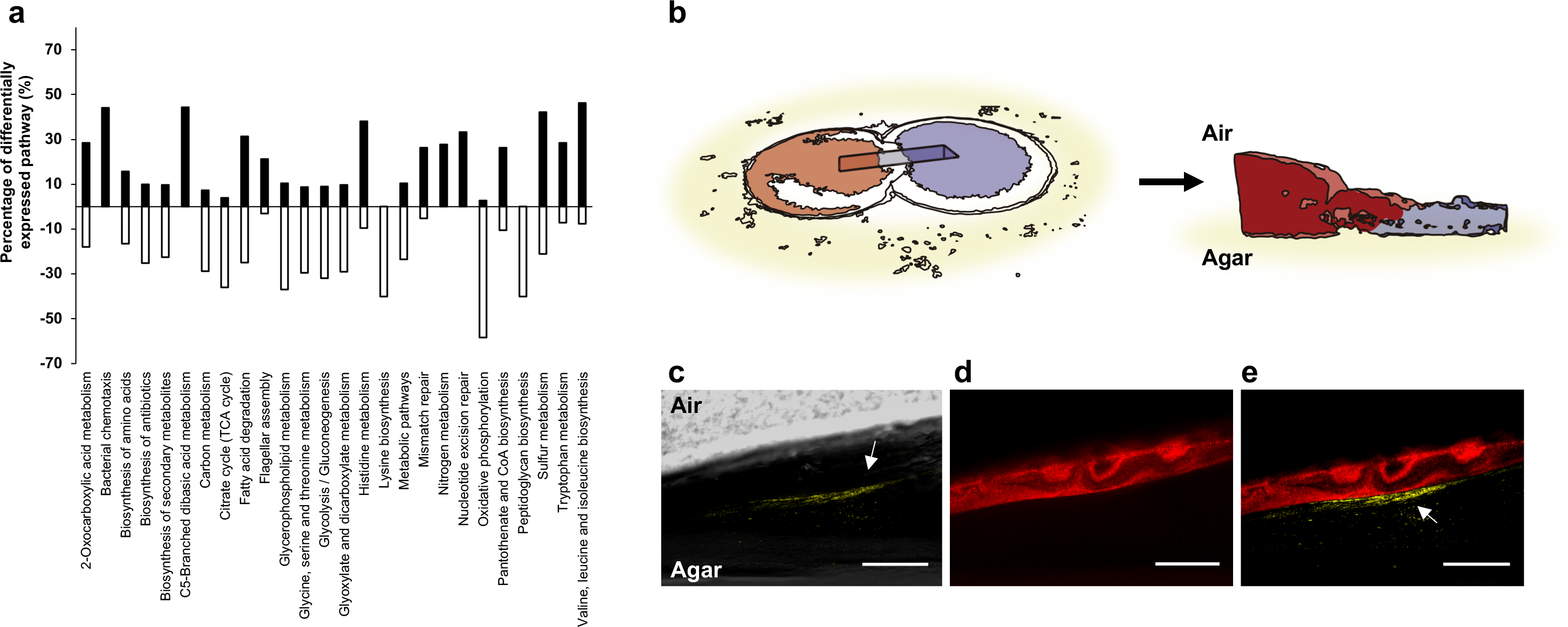
Non-sporulated *Bacillus* Δmatrix cells activate chemotaxis and motility in response to the PCL1606 colonization. (a) KEGG pathway analysis of the Δmatrix genes induced (black bars) and repressed (empty bars) after 72 h of interaction with PCL1606 (*P*-value < 0.05). (b, c, d and e) Confocal laser scanning microscopy (CLSM) images of cross-sections of the interaction between PCL1606 (DsRed) and Δmatrix *P*_*mot*_- YFP after 72 h of growth. (b) An illustration of the interaction PCL1606 (red) and Δmatrix (blue) and the cross-section analyzed in CLSM. (c) Fluorescence from the YFP channel within the Δmatrix *P*_*mot*_-YFP cells is shown in yellow. (d) Fluorescence from the RFP channel within the PCL1606 cells is shown in red. (e) Merged channels image of the interaction Δmatrix *P*_*mot*_-YFP and PCL1606. Scale bar, 100 μm.

To assess this idea, the promoter of *mot*, a gene upregulated in the interaction of Δmatrix with PCL1606, was fused to YFP and the cellular pattern of expression examined at 72 h using fluorescence microscopy (Fig. 6b-e). Most of the Δmatrix population did not express detectable levels of fluorescence (Fig. 6c), which is consistent with our previous observations indicating that most of the Δmatrix population had sporulated at this point. However, we could detect some regions in close proximity to PCL1606 cells (Fig. 6d) where *P*_*mot*_ was active and expressing detectable amounts of YFP. This suggests that the *mot* promoter is indeed active in a small percentage of the *Bacillus* population and may reflect vegetative bacteria responding to the presence of PCL1606 cells by increasing their motility (Fig. 6e).

### The two distinctive strategies of *Bacillus* and *Pseudomonas* permit their coexistence on plants

Our *in vitro* experiments have shown that *Pseudomonas* uses the T6SS as a powerful offensive strategy to exclude or compete with competitors, and that *Bacillus* responds to this behavior with a well-structured extracellular matrix and sporulation as a second line of a defensive strategy. To better understand the ecological significance of these two strategies, we examined these interactions on two anatomically and chemically different melon plant organs: the leaves, where bacterial cells have to adapt to limited nutrients and space (Fig. 7), and the seeds, where germination give rises to an emergence of nutrients and a nascent radicle susceptible for colonization where bacterial species are in close contact (Fig. 8).

**Fig. 7.**
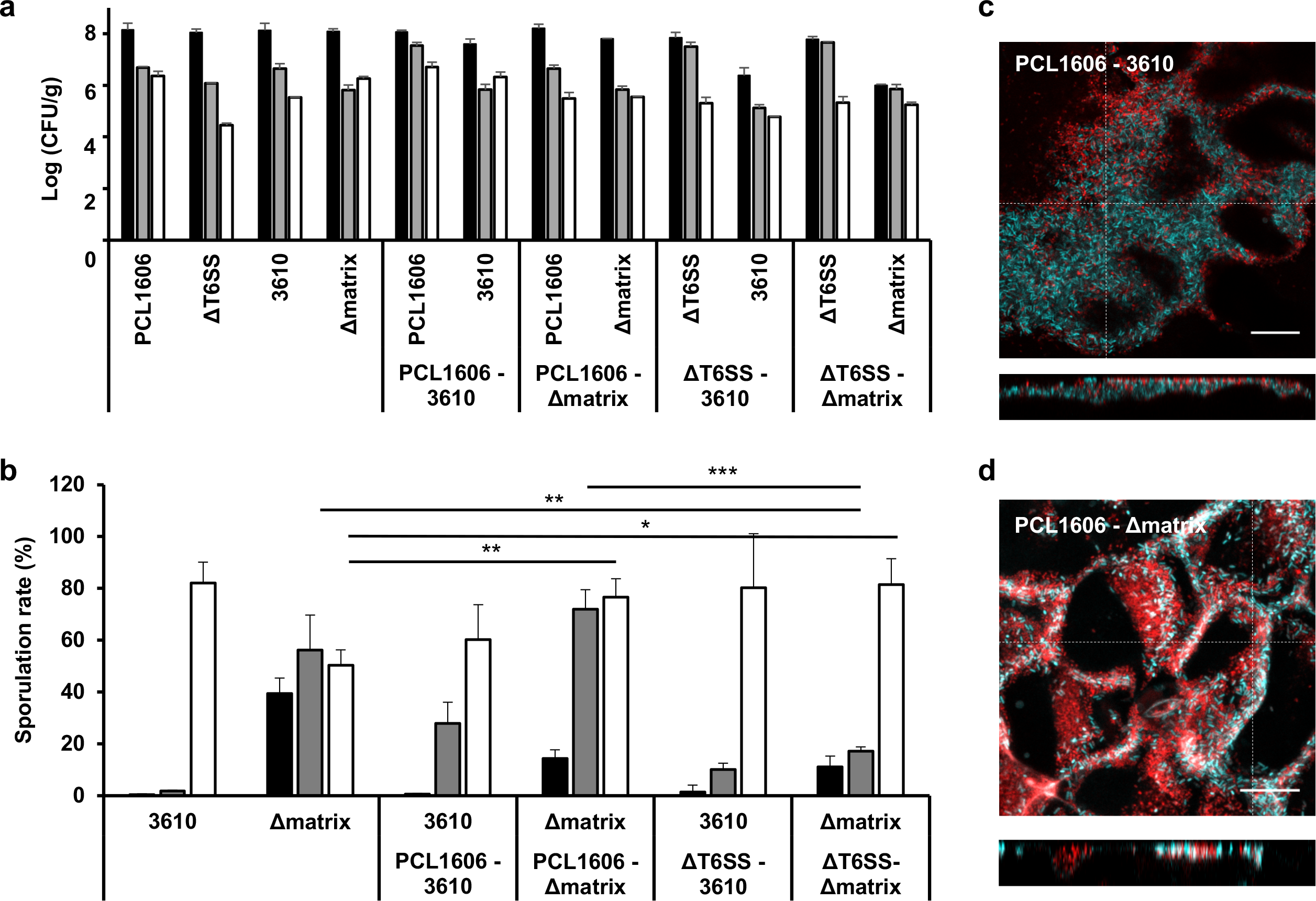
The extracellular matrix and sporulation of *Bacillus* permits the coexistence with *Pseudomonas* on plant leaves. (a) Log (CFU/ml) of *B. subtilis* (3610 and Δmatrix) and *P. chlororaphis* (PCL1606 and ΔT6SS) strains after single- and co-inoculation of melon leaves. (b) Sporulation percentages of *Bacillus subtilis* species after single- and co-inoculation of melon leaves. Colony forming units and spore percentages were counted at 0 dpi (black bars), 2 dpi (grey bars) and 9 dpi (white bars). Average values of five biological replicates are shown, with error bars representing SD. \**P*value <0.05, ** *P* value <0.01, \*\*\**P* value <0.001 (Tukey test). (c) CLSM maximum projections and z axis slices at positions indicated by the discontinuous lines of the interactions between 3610 - PCL1606 (top) and Δmatrix – PCL1606 (bottom) after 9 days of co-inoculation. *Bacillus* strains were labeled with CFP and PCL1606 was labeled with DsRed.

**Fig. 8.**
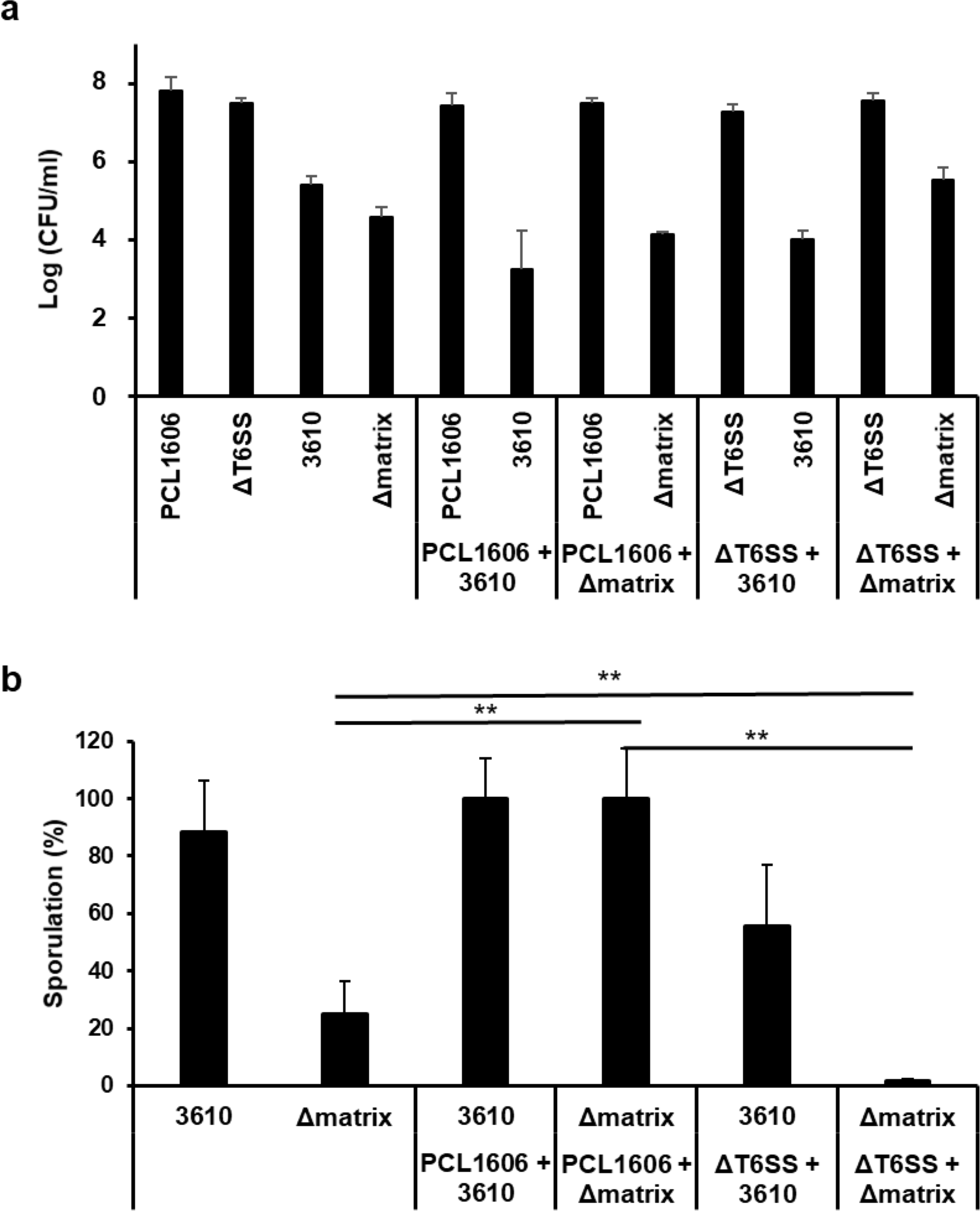
The T6SS of *Pseudomonas* kills and triggers the sporulation of *Bacillus* in melon seed radicles. (a) Log (CFU/ml) of *B. subtilis* (3610 and Δmatrix) and *P. chlororaphis* (PCL1606 and ΔT6SS) strains after single- and co-bacterization of melon seeds. (b) Sporulation percentages of *Bacillus subtilis* strains after single- or co-inoculation with *Pseudomonas* strains of melon seeds. Colony forming units and spore percentages of *Bacillus* were counted 5 dpi. Average values of four biological replicates are shown, with error bars representing SD. \*\**P* value <0.01 (Tukey test).

We examined the persistence (CFU/ml) and colonization patterns of the different strains by CLSM analysis at three time-points during the nine days after bacterial inoculation on melon leaves. The persistence patterns of both *Bacillus* strains in melon leaves were similar in all the cases and regardless single or co-inoculations, with small decreases in the bacterial populations after 2 or 9 days of inoculation (Fig. 7a). CLSM images showed co-localization of the two species but with a stratified arrangement of PCL1606 and 3610 WT strains mostly in the intercellular zones of surface plant cells (Fig. 7c). Co-inoculation of Δmatrix and PCL1606 led to a more dispersed and heterogeneous distribution of cells lacking the horizontal stratification seen with 3610 (Fig. 7d). Quantification of bacterial distribution relative to leaf surfaces confirmed this change in organization with the Δmatrix population (Suppl. Fig. 11b and d) being displaced further from the leaf surface versus the wild-type strain (Suppl. Fig. 11a and c) in agreement with a role for the *Bacillus* extracellular matrix as a physical barrier or support.

The higher levels of sporulation of both *Bacillus* strains in co-inoculated leaves compared to mono-inoculated ones suggests that the sporulation survival strategy observed *in vitro* is also active in more complex real-world environments such as leaves (Fig. 7b). However, in contrast to our *in vitro* observations, when assays were performed using ΔT6SS instead of the WT PCL1606 strain, a noticeable delay in the *Bacillus* sporulation rate was observed, although sporulation reached similar levels to those obtained in mono-inoculated leaves after 9 days, suggesting the involvement of the T6SS in the triggering of the *Bacillus* sporulation in plant leaves (Fig. 7b).

The same co-inoculation experiments were done in melon seeds, as both strains are known to promote seed germination and to live in association with plant rhizospheres. The CFU counts after 5 days of seed germination revealed a reduction of two orders of magnitude of the 3610 population when co-cultured with PCL1606 (Fig. 8a). Furthermore, as previously seen in melon leaves, the 3610 population had almost completely sporulated after 5 days alone or co-inoculated with PCL1606. However, the Δmatrix population showed remarkable sensitivity to the presence of PCL1606, with a sporulation rate rising from 27% in the mono-bacterized seeds to close to 100% in the co-bacterized samples (Fig. 8b). Interestingly, the Δmatrix population increased in one order of magnitude when co-inoculated with PCL1606 ΔT6SS compared to the single inoculated (Fig. 8a). In addition, the co-bacterization of 3610 or Δmatrix with the ΔT6SS resulted in the reduction of the sporulation rate, becoming almost negligible in Δmatrix (Fig. 8b).

These observations demonstrate that bacterial interactions are mediated by common mechanisms in vastly different environments. Although the relative importance of these mechanisms may vary in different environments, we have shown that the *Bacillus* extracellular matrix and the *Pseudomonas* T6SS play important roles in establishing the balance between different bacterial populations. Further studies into the roles of other factors identified in this study, such as surfactin, may help uncover other ways in which bacteria defend themselves and even co-operate with other bacterial species.

## Discussion

Understanding the behavior and mechanisms involved in the interaction between bacterial species that share ecological niches is fundamental in the study of microbial population evolution and has important implications for biotechnology^41^. In their interaction with plants, it is important to decipher how intrinsically diverse microbes are related to each other, the potential strategies used in host adaption and how these strategies might coexist to produce a microbial consortium that benefits the host plant^42^. Many of the factors that affect microbial interactions are mediated via cell-to-cell contact or by diffusible molecules. Therefore, cellular spatial distribution is a key environmental factor in determining the result of these microbial interactions^43^. This suggests important role for biofilms, which provide protection against harsh environments, enhance tolerance to physical and chemical stresses, and promote metabolic cooperation and the community-coordinated adjustment of gene expression^44, 45^. In this work we provide unprecedented insights into the fundamentals that underlie social inter-species interactions by studying how *Bacillus subtilis* NCIB3610 and *Pseudomonas chlororaphis* PCL1606 interact and coexist *in vitro* and on plants.

*In vitro* experiments in optimal media for each bacterial species resulted in the physical exclusion of each species, and most importantly, the reduction of wrinkles, the most noticeable morphological feature of *B. subtilis* biofilms (Fig. 1a and d). Based on our examination of pairwise interactions we have confirmed that the main components of the *Bacillus* extracellular matrix (Eps, TasA and BslA) play a critical role in the protection of the *Bacillus* colony from the invasiveness of *Pseudomonas*. More specifically, we can assign to this defensive strategy major contributions by the exopolysaccharide and BslA, the two main factors contributing to the hydrophobicity and water repellence of *Bacillus* biofilms and its resistance to water dissolved and diffusible active molecules^16, 20^. However, the fact that both wild-type and the extracellular matrix-deprived Δmatrix strain, sporulated massively following contact with *Pseudomonas* suggests that some molecules are able to diffuse through this polymeric armor and trigger sporulation. Thus, it is reasonable to think that the extracellular matrix might be physically impeding access of *Pseudomonas* cells into the *Bacillus* colony, a notion supported by the overgrowth of *Pseudomonas* on the Δmatrix colony (Fig. 1e). These findings highlight and support the complexity of the extracellular matrix of *Bacillus* and the complementary contribution of each structural component to the full functionality of this megastructure.

Examination of this interaction under the microscope allowed us to delineate more precisely the sequence of events at earlier stages, before macroscopic changes are visible, revealing details not observed in previous studies. First, the absence of extracellular matrix permitted faster expansion of the *Bacillus* colony, leading to the Δmatrix strain making contact with PCL1606 earlier than the wild-type strain, and the partial penetration of *Pseudomonas* by the Δmatrix colony. Surfactin is a potent surfactant that reduces the water surface tension and promotes the social movement of *B. subtilis*^49, 50^. Up to now, bacterial interactions involving surfactin-producing *Bacillus* species have suggested an inhibitory role for surfactin against other *Bacillus* species, or as an inhibitor of *Streptomyces* aerial hyphae development^51^. However, in this work we have presented evidence that suggests a novel role for surfactin. Imaging mass spectrometry analysis suggests an accumulation of surfactin mostly in the zone of the initial contact and a distribution pattern altered in the Δmatrix strain. In contrast to the function described for interactions with other *Bacillus* species where surfactin showed an antimicrobial effect^46^, in this case surfactin appears to collaterally promote *Pseudomonas* colony spread. Therefore, we propose surfactin as a common mechanism for promoting interspecies interaction, with further progression of the interaction, positive or negative, influenced by additional factors specifically employed by each bacterial species^37, 52^.

Further evaluation of the Δmatrix-PCL1606 interaction revealed the overgrowth phenotype of *Pseudomonas* to be associated in part with its ability to metabolically adapt to the new environment found in the Δmatrix area of the interaction. An additional structural change in PCL1606 upon interaction with *Bacillus* was activation of the T6SS. The T6SS encoded in the PCL1606 genome is genetically similar to the H2-T6SS of *Pseudomonas aeruginosa*^29^ but with the peculiarity of having a PAAR gene and two genes of unknown functions in the middle of the T6SS cluster. Up to now, many works have shown a role for T6SS against Gram-negative bacteria species and eukaryotic cells^53, 54, 55^, but no studies have reported the efficiency of the T6SS against Gram-positive bacteria. In addition, recent work has highlighted the relevance of exopolysaccharide from *Vibrio cholerae* in the protection against external T6SS^56^. However, this is the first demonstration that the T6SS can also attack Gram-positive bacteria. Our findings indicate that the immunity of *Bacillus* to this offensive tool of *Pseudomonas* seems to be depended on the synthesis of an extracellular matrix, with the activation of sporulation as a secondary defense strategy in response to the *Pseudomonas* T6SS, and the maintaining of a small sub-population able to move and escape from the *Pseudomonas* presence by the activation of motility elements.

Based on our studies, we propose two mechanisms that induce *Bacillus* sporulation depending on the distance between cells. The first mechanism would be inflicted by diffusible molecules produced by *Pseudomonas*, and with no need of cellular contact, as supported by the sporulation of both 3610 WT and Δmatrix prior to direct contact with PCL1606. Previous reports have proven that siderophores^57^ and compounds such as decoynine and hadacidin^58^ promote the sporulation of *Bacillus*. *P. chlororaphis* PCL1606 produces pyochelin, a potent siderophore that might retain the same functionality, however, neither transcriptomic or MALDI-IMS analyses revealed differences in the transcriptional level or distribution pattern of this molecule in the different pairwise interactions. Moreover, purified pyochelin failed to activate expression from the sporulation-related *sspB* promoter or change the rate of *B. subtilis* sporulation (Data not shown). Similar results were obtained with different fractions purified from *P. chlororaphis* cultures. Thus, we suspect that the induction of *Bacillus* sporulation is most likely multifactorial. The other strategy is most likely mediated by close contact between *B. subtilis* and *P. chlororaphis* cells, and we propose the T6SS as the additional trigger for *Bacillus* sporulation as observed in the plant experiments.

The results from the plant experiments support the value of the mechanistic findings of the interactions evaluated *in vitro*, and highlight the importance of the physical distance of cells in defining the final result of the interactions. Our findings highlight the relevance of the extracellular matrix of *B. subtilis* for surviving, persisting and colonizing two different plant organs^4^, and the role of sporulation as a secondary defensive strategy in the close contact with *Pseudomonas* cells. The T6SS of *P. chlororaphis* seems to have adopted a double role in these scenarios: i) as a killing factor, reducing the cell density of *Bacillus* in the radicle, and ii) as a trigger of sporulation, delaying or reducing the sporulation rates of *Bacillus* strains in leaves and melon radicles when a *Pseudomonas* ΔT6SS strain is used (Fig. 7b and 8b). In summary, our work increases our understanding of the mechanics of the complex and multifactorial bacterial social interactions, demonstrating that the adaptive strategies adopted by two bacterial species may coexist and lead to the formation of stable bacterial communities in plants, either mixed or as physically separated sub-domains. The findings in this work deepen our understanding of how mixed microbial communities might have evolved and provide valuable knowledge that may aid in the manipulation of bacterial consortium for agricultural and other applications.

## Materials and Methods

### Strains, media and culture conditions

A complete list of the bacterial strains used in this study is shown in Suppl. Table 3. Routinely, bacterial cells were grown in liquid LB medium at 30°C (*P. chlororaphis* and *B. subtilis*) or 37°C (*E. coli*) with shaking on an orbital platform. When necessary, antibiotics were added to the media at appropriate concentrations. Strains and plasmids were constructed using standard methods^59^. Oligonucleotides used in this study are listed in Suppl. Table 4.

### *Pseudomonas* T6SS mutant

Chromosomal deletion of ImpA (TssA), a core T6SS baseplate component essential for its activity^60^, was performed using the I-SceI methodology^61, 62, 63^ in which upstream and downstream segments of homologous DNA are separately amplified and then joined to a previously digested pEMG vector using the Gibson Assembly Master Mix^64^. The resulting plasmid was then electroporated into *P. chlororaphis* PCL1606. After selection for positive clones, the pSEVA628S I-SceI expression plasmid was also electroporated. Cointegrated constructions were resolved by induction of I-SceI expression with 3 mM 3-methylbenzoate. Kanamycin-sensitive clones were PCR analyzed to verify the deletions. The pSEVA628S plasmid was cured by growth without selective pressure and its loss confirmed by sensitivity to 30 μg ml^−1^ gentamicin and colony PCR screening as described by Martinez-Garcia and de Lorenzo ^62^.

### *Bacillus subtilis* extracellular matrix mutants

*Bacillus subtilis* mutants were generated by SPP1 phage transduction as previously described^65^. To obtain the double mutant strains, phage lysates from *tasA* and *eps* single mutant strains were obtained and transferred to a *bslA* mutant. The same procedure was used to obtain the triple mutant, in this case using lysates from the *eps* single mutant and transferring it to a *bslA*-*tasA* double mutant. Mutants were confirmed both by PCR and by antibiotic resistance (kanamycin, MLS and tetracycline in the case of the matrix mutant strain).

### Construction of fluorescence labeling strains

Fluorescence labeling plasmid pKM008V was constructed for *B. subtilis* strains. Briefly, the P_veg_ promoter fragment (300 bp) was extracted from pBS1C3 by digestion with EcoRI and HindIII restriction enzymes, purified and cloned into pKM008 plasmid, which was previously digested with the same restriction enzymes. We used P_veg_ as it is considered a constitutive promoter in *Bacillus subtilis*. The same procedure was followed for mot promoter but the fragment was obtained by PCR using the 3610 chromosome as template. pKM008V was then transformed into *B. subtilis* 168 by natural competence, and transformants selected by plating in LB plates supplemented with spectinomycin (100 μg ml^−1^). Finally, the extracellular matrix mutant was fluorescently marked by transferring CFP from *B. subtilis* 168 using SPP1 phage transduction as previously described^65^.

In the case of *P. chlororaphis* PCL1606 P_T6SS_-DsRed, we fluorescently labeled the PCL1606 strain using pSEVA237D. Briefly, we amplified the P_T6SS_ promoter region (250 bp) from PCL1606 genomic DNA. Plasmid and fragment were digested with EcoRI and HindIII restriction enzymes, purified, ligated and cloned in *E. coli* DH5α competent cells. The completed pSEVA237D-P_T6SS_ was then electroporated into *P. chlororaphis* PCL1606 cells. Introduction of the plasmid was confirmed by PCR and antibiotic selection (kanamycin).

### Pairwise interactions

*Bacillus subtilis* and *Pseudomonas chlororaphis* strains were routinely spotted at a 0.7 cm distance onto an LB agar plate using 2 μl of cell suspension at an OD600 of 0.5. Plates were incubated at 30°C and images taken at different time points. Photographs were captured using a Panasonic DMC-FZ250 camera. For confocal microscopy time-course experiments, 0.7 μl of cell suspension were spotted at a 0.5 cm distance onto 1.2 mm thick LB agar plates using 35mm glass bottomed dishes suitable for confocal microscopy (Ibidi).

### Bacterial population dynamics

To analyze the cell-number percentage and growth curves during the interaction between *B. subtilis* (3610 and Δmatrix) and *P. chlororaphis* PCL1606 (PCL1606 and ΔT6SS), interaction plates were prepared as described, and incubated for the required time period. The interaction was then divided into three sections. The *B. subtilis* section (*Bacillus* Area) consisted of the entire *B. subtilis* colony, not including the area proximal to the *P. chlororaphis* colony. The interaction section (Intermediate Area) consisted of *B. subtilis* and *P. chlororaphis* cells more proximal to the other colony. The *P. chlororaphis* section (*Pseudomonas* Area) consisted on the entire PCL1606 colony excluding the cells included in the intermediate area. Each section was inserted into a 1.5 ml eppendorf tubes containing 1 ml PBS, completely resuspended, diluted serially and plated onto LB petri dishes. Plates were incubated overnight at 30°C. *Pseudomonas* and *Bacillus* colonies were easy to differentiate in terms of counts as they are morphologically different. For spores count, the serial dilutions previously mentioned were heated to 80°C for 10 minutes in order to kill non-sporulated bacteria and plated as mentioned above.

### Whole-genome transcriptomic analysis

Single colonies of *P. chlororaphis* PCL1606 and *B. subtilis* NCIB3610 (Δmatrix) were grown overnight in LB medium at 30°C and spotted as single colonies or as interactions as previously described for 72 h. After that, cells were collected and stored at −80°C. All the assays were performed in duplicate. Single colonies (control) were collected completely, while in pairwise interactions we only collected the area where *Bacillus* and *Pseudomonas* were mixed. For disruption of *Bacillus* single colonies and *Pseudomonas-Bacillus* mixed colonies, collected cells were resuspended in BirnBoim A (Birnboim and Doly 1979) and lysozyme was added and incubated 15 min at 37°C. After that, suspensions were centrifuged, the pellet resuspended in Trizol and total RNA extraction performed as indicated by the manufacturer. RNA extraction of *Pseudomonas* control colonies started in this point of the protocol as it is not necessary to add lysozyme for cell disruption. DNA removal was carried out by treatment with Nucleo-Spin RNA Plant (Macherey-Nagel). Integrity and quality of total RNA was assessed with an Agilent 2100 Bioanalyzer (Agilent Technologies). Removal of rRNAs was performed using RiboZero rRNA removal (bacteria) kit from Illumina and 100-bp single-end reads libraries were prepared using a TruSeq Stranded Total RNA Kit (Illumina). Libraries were sequenced using a NextSeq550 sequencer (Illumina).

Raw reads were quality-trimmed with cutadapt. Then, trimmed reads were mapped against *Pseudomonas chlororaphis* PCL1606, *Bacillus subtilis* 3610 reference genomes using EDGE-pro^66^ software, which is specially designed to process prokaryotic RNA-Seq data.

Quantification was also performed with EDGE-pro. Raw counts were normalized using the trimmed mean of M-values (TMM) method^67^, implemented in NOISeq R package^68^. Differential expression analyses were performed with DESeq2 package^69^. Genes were considered as differentially expressed when logFC was higher than 1 and lower than −1, and p-value < 0.05. Data have been deposited in the GEO database: GSE117802.

### Confocal Laser Scanning Microscopy

Bacterial interactions in solid medium were visualized by Confocal Laser Scanning Microscopy (CLSM). For the interaction between Δmatrix labeled with CFP and *P. chlororaphis* PCL1606 PT6SS-DsRed and Δmatrix P_mot_-YFP versus PCL1606, single colonies and bacteria growing in direct contact were grown on LB solidified with 1.5% agar as described above. A quarter section of the colony (after 72h of growth) was excised with a surgical scalpel and placed in a transversal position directly onto a 24 mm diameter round 1.5 thickness coverslip using a 35mm coverslip adaptor suitable for inverted microscopes. For the observation of *B. subtilis* and *P. chlororaphis* strains (labelled with CFP and DsRed, respectively) growing on melon leaves, 30 mm diameter discs were obtained with a cork-borer. A drop of glycerol was applied onto a labeled section each colony and placed onto 1.5 thickness cover glass (22 × 22 mm) and sealed with adhesive tape to a standard glass microscope slide. In both cases, images were obtained by visualizing samples using a Leica SP5 confocal microscope with a 40× NA 1.3 Plan APO oil immersion objective. Image processing was performed using Leica LAS AF (LCS Lite, Leica Microsystems) and FIJI/ImageJ software. For each experiment, laser settings, scan speed, PMT detector gain, and pinhole aperture were kept constant for all acquired image stacks.

For bacterial interaction time-course experiments, *B. subtilis* and *Pseudomonas chlororaphis* labeled strains were placed on a thin (1.2 mm) layer of LB agar in 35mm glass bottom dishes (Ibidi) suitable for microscopy. Plates were incubated at 30°C for 6 h prior to acquisition. Temperature was maintained at 30°C during the time-course using the integrated microscope incubator. Acquisitions were performed using an inverted Leica SP5 confocal microscope with a 25× NA 0.95 NA IR APO long working distance water immersion objective. Bacterial fluorescence could be visualized from underneath the bottom of the plate and through the agar medium thanks to the long 2.2 mm free working distance of this objective. A special oil immersion medium, Immersol W 2010 (Zeiss) was used instead of water to avoid problems with evaporation during the experiment. Colony fluorescence was followed in multiple regions selected at the start of the experiment, with the acquisition of a series of different focal (z) positions at each region performed automatically at every time-point. Evaporation from the LB agar and its utilization by the growing colonies resulting a gradual lowering of the agar surface relative to the objective lens of approximately 250 microns every 24 hours. In order to be able to follow colony dynamics, images were acquired over wide focal range to compensate for the predicted change in colony position during the experiment. Image processing and 3D visualization was performed using ImageJ/FIJI^70, 71^ and Imaris version 7.6 (Bitplane).

Colony growth speeds were calculated with FIJI, using the change in position of the leading edge of the colony between time-points to calculate its speed in microns per seconds. Results were calculated as an average speed at three different regions of the same colony, with variations expressed as standard deviation. To reduce the impact of random vibrations and variations, plotting speed values were smoothed as a floating 4-value average advancing 30 minutes (or 1 timepoint) at a time.

### Matrix-assisted laser desorption ionization imaging mass spectrometry (MALDI-IMS)

To perform MALDI-IMS a small section of LB agar containing the cultured microorganisms (both in single colonies and in interactions) were cut and transferred to a MALDI MSP 96 anchor plate. Deposition of universal matrix (1:1 mixture of 2,5-dihydroxybenzoic acid and α-cyano-4-hydroxy-cinnamic acid) over the agar was done using a 53 μm molecular sieve. After that, plates were dried at 37 °C for 4 hours. Photographs were taken before and after matrix deposition. Samples were analyzed using a Bruker Microflex MALDI-TOF mass spectrometer (Bruker Daltonics, Billerica, MA, USA) in positive reflectron mode, with 300 μm−400 μm laser intervals in X and Y directions, and a mass range of 100-3200 Da. Data obtained were analyzed using FlexImaging 3.0 software (Bruker Daltonics, Billerica, MA, USA).

### Liquid chromatography - tandem Mass Spectrometry (LC-MS/MS)

Non-targeted LC-MS/MS analysis was performed on a Q-Exactive Quadrupole-Orbitrap mass spectrometer coupled to Vanquish ultra-high performance liquid chromatography (UPLC) system (Thermo Fisher Scientific, Bremen, Germany) according to Petras, Nothias 72 Therefore, 5 μL of the samples were injected on UHPLC separation on C18 core-shell column (Kinetex, 50 × 2 mm, 1.8 um particle size, 100 A pore size, Phenomenex, Torrance, USA). For the mobile phase we used a flow rate of 0.5 mL/min (Solvent A: H2O+ 0.1 % formic acid (FA), Solvent B: Acetonitrile (ACN) + 0.1 % FA). During the chromatographic separation, we applied a linear gradient from 0-0.5 min, 5 % B, 0.5-4 min 5-50 % B, 4-5 min 50-99 % B, flowed by a 2 min washout phase at 99% B and a 2 min re-equilibration phase at 5 % B. For positive mode MS/MS acquisition the electrospray ionization (ESI) was set to a 35 L/min sheath gas flow, 10 L/min auxiliary gas flow, 2 L/min sweep gas flow and 400 °C auxiliary gas temperature. The spray voltage was set to 3.5 kV with an inlet capillary of 250 °C. The S-lens voltage was set to 50 V. MS/MS product ion spectra were acquired in data dependent acquisition (DDA) mode. MS1 survey scans (150-1500 m/z) and up to 5 MS/MS scans per DDA duty cycle were measured with a resolution (R) of 17,500. The C-trap fill time was set to a maximum of 100 ms or till the AGC target of 5E5 iones was reached. The quadrupole precursor selection width was set to 1 m/z. Normalized collision energy was applied stepwise at 20, 30 and 40 % with z = 1 as default charge state. MS/MS scans were triggered with apex mode within 2 to 15 s from their first occurrence in a survey scan. Dynamic precursor exclusion was set to 5s. Precursor ions with unassigned charge states and isotope peaks were excluded from MS/MS acquisition.

### Data Analysis and MS/MS Network Analysis

After LC-MS/MS acquisition, raw spectra were converted to.mzXML files using MSconvert (ProteoWizard). MS1 and MS/MS feature extraction was performed with Mzmine2.30^73^. For MS1 spectra an intensity threshold of 1E5 and for MS/MS spectra of 1E3 was used. For MS1 chromatogram building a 10 ppm mass accuracy and a minimum peak intensity of 5E5 was set. Extracted Ion Chromatograms (XICs) were deconvoluted using the baseline cut-off algorithm at an intensity of 1E5. After chromatographic deconvolutiuon, XICs were matched to MS/MS spectra within 0.02 m/z and 0.2 min retention time windows. Isotope peaks were grouped and features from different samples were aligned with 10 ppm mass tolerance and 0.1 min retention time tolerance. MS1 features without MS2 features assigned were filtered out the resulting matrix as well as features which did not contain isotope peaks and which did not occur at least in 3 samples. After filtering gaps in the feature matrix were filled with relaxed retention time tolerance at 0.2 min but also 10 ppm mass tolerance. Finally, the feature table was exported as.csv file and corresponding MS/MS spectra as.mgf file. Contaminate features observed in Blank samples were filtered and only those whose relative abundance ratio blank to average in the samples was lower than 50% were considered for further analysis.

For molecular networking and spectrum library matching the.mgf file was uploaded to GNPS (gnps.ucsd.edu)^74^. For molecular networking, the minimum cosine score was set as 0.7 The Precursor Ion Mass Tolerance was set to 0.01 Da and Fragment Ion Mass Tolerance to 0.01 Da, Minimum Matched Fragment Peaks was set to 4, Minimum Cluster Size to 1 (MS Cluster off) and Library Search Minimum Matched Fragment Peaks to 4. When Analog Search was performed the Cosine Score Threshold was 0.7 and Maximum Analog Search Mass Difference was 100. Molecular networks were visualized with Cytoscape version 3.4^75^.

### MS/MS Data availability

All LC-MS/MS data was deposited to the Mass spectrometry Interactive Virtual Environment (MassIVE) at https://massive.ucsd.edu/ with the identifier MSV000082402. Molecular Networking and spectrum library matching results can be found online at the GNPS webpage under the following links: https://gnps.ucsd.edu/ProteoSAFe/status.jsp?task=705915a36bd24cce9dbcedd4b876b008 and https://gnps.ucsd.edu/ProteoSAFe/status.jsp?task=2c41ab80bb12490ba52fa7e8c08fd56b.

### Bacterial interactions on plant leaves

The assay was set up as previously reported ^76^. Briefly, melon seeds of cv. Rochet were pre-germinated, sown into pots and cultivated in a plant growth chamber until use. Before each experiment, bacterial cultures were incubated overnight at 28°C in an orbital shaker, two-time washed and cultures adjusted to a cell density of 10^8^ cfu ml^−1^. In the case of *Bacillus* and *Pseudomonas* co-inoculation, cultures were mixed prior to leaf application. Plants were then incubated in a growth chamber under controlled conditions (25°C over a 16-8 h photoperiod). Leaves were collected at 4 hours, 2 days and 9 days post-inoculation, fresh weight was measured and CFU and spore percentages calculated. 3D acquisitions of melon leaf surfaces were acquired with a 40× 1.30 NA Plan APO oil immersion objective and Leica SP5 confocal microscope. CFP-positive 3610 and DsRed-positive 1606 bacteria were detected automatically using the Imaris (version 7.6.5) spot detection algorithm using an estimated diameter of 1.26 μM and background subtraction. Identical CFP and DsRed intensity thresholds were used between samples. Leaf surfaces were manually defined as an Imaris surface object and used to calculate the number of CFP and DsRed positive bacteria at distances of 1 to 9 μM from the leaf surface at 1 micron intervals using the Imaris “Find Spots Close to Surface” function. At all-time points, leaf discs were taken for confocal microscopy analysis. Experiments were repeated at least three times.

### Seed colonization assays

Melon seeds were bacterized with mono- and co-cultures of *B. subtilis* and *P. chlororaphis* for one hour. Competitive colonization assays were performed as previously described^77^, using single strains and 1:1 mixes of *B. subtilis* and PCL1606 strains. Seeds were grown for 5 days at 25°C before bacterial persistence quantification and size, weight and area calculation. For bacterial quantitative analysis, roots were cut, weighed and introduced in eppendorfs with 1 ml of M9 basal medium and 1 g of glass beads (diameter 3 mm). Bacteria attached to the roots were recovered by vortexing for 1 min, and colony forming units and sporulation percentages were calculated by plating serial dilutions of the resulting suspension on LB medium. Antibiotics supplementation was not needed as colonies were easily differentiated by morphology. Experiments were repeated at least three times.

### Statistical analysis

Results are expressed as mean ± SD. Statistical significance was assessed using Tukey’s test. All analyses were performed using GraphPad Prism^®^ version 6. P-values <0.05 were considered significant.

## Acknowledgments

We thank the Ultrasequencing Unit of the SCBI-UMA for RNA sequencing, and the Bioinformatic unit of GENYO (Granada, Spain) for the analytical treatment of the data. We are grateful to Marta Martínez-Gil (University of Málaga) and Paul Straight (Texas A&M University) for critical reading and multiple suggestions and comments on the manuscript, and Luis Díaz (University of Málaga) for technical support in some illustrations. This work was partially supported by grants from ERC Starting Grant (BacBio 637971) and Plan Nacional de I+D+I of Ministerio de Economía y Competitividad (AGL2016-78662-R). C.M.S is funded by the program Juan de la Cierva Formación (FJCI-2015-23810). M.V.B.C is funded by the program Plan Propio de Investigación y Transferencia from Universidad de Málaga. Furthermore, we would like to thank Ministerio Economía y Competitividad for their support through grant AGL2014-5218-C2-1-R to F.M.C., and the US National Institutes of Health (NIH) for their support through grants GMS10RR029121, 5P41GM103484-07 to P.C.D. and the German Research Foundation (DFG) with Grant PE 2600/1 to D.P.

## Author contributions

D.R conceived the study; D.R. and C.M.S designed the experiments; C.M.S. performed the main experimental work; C.M.S and J.P performed and designed the confocal microscopy work and data analysis; Y.N.G constructed some *Bacillus* strains, C.M.S. and M.V.B.C performed the experimental work related with seeds; C.M.S, D.P and A.M.C.R performed MALDI-IMS and LC-MS/MS analysis; D.R and C.M.S wrote the manuscript; and J.P, A.V, F.M.C., and P.C.D contributed critically to writing the final version of the manuscript.

